# Influenza virus replication in cardiomyocytes drives heart dysfunction and fibrosis

**DOI:** 10.1101/2021.10.27.466128

**Authors:** Adam D. Kenney, Stephanie L. Aron, Clara Gilbert, Naresh Kumar, Peng Chen, Adrian Eddy, Lizhi Zhang, Ashley Zani, Nahara Vargas-Maldonado, Samuel Speaks, Jeffrey Kawahara, Parker J. Denz, Lisa Dorn, Federica Accornero, Jianjie Ma, Hua Zhu, Murugesan V.S. Rajaram, Chuanxi Cai, Ryan A. Langlois, Jacob S. Yount

**Affiliations:** Department of Microbial Infection and Immunity, The Ohio State University, Columbus, OH, USA; Infectious Diseases Institute, Viruses and Emerging Pathogens Program, The Ohio State University, Columbus, OH, USA; Department of Microbiology and Immunology, The University of Minnesota, Minneapolis, MN, USA; Department of Physiology and Cell Biology, The Ohio State University, Columbus, OH, USA; Department of Surgery, The Ohio State University, Columbus, OH, USA

## Abstract

Cardiac dysfunction is a common extrapulmonary complication of severe influenza virus infection. Prevailing models propose that influenza-associated heart dysfunction is indirectly triggered by cytokine mediated cardiotoxicity downstream of the inflamed lung, rather than by direct infection of cardiac tissue. To test the etiology of cardiac dysfunction resulting from influenza virus infection, we generated a novel recombinant H1N1 influenza A virus that was attenuated in cardiomyocytes by incorporation of target sequences for miRNAs expressed specifically in that cell type (miR133b and miR206). Compared with control virus, mice infected with the miR-targeted virus had significantly reduced heart viral titers, confirming cardiac attenuation of viral replication. The miR-targeted virus, however, was fully replicative and inflammatory in lungs when compared to control virus, and induced similar systemic weight loss. The miR-targeted virus induced considerably lower levels of cardiac arrhythmia, fibrosis, and inflammation, compared with control virus, in mice lacking interferon induced transmembrane protein 3 (IFITM3), which serve as the only available model for severe influenza-associated cardiac pathology. We conclude that robust replication of virus in the heart is required for pathology even when lung inflammation is severe. Indeed, we show that human stem cell-derived cardiomyocytes are susceptible to influenza virus infection. This work establishes a fundamental new paradigm in which influenza virus damages the heart through direct infection of cardiomyocytes.

## Introduction

Seasonal influenza virus remains a major contributor to human mortality, and the potential for emergence of new pandemic strains is an ever-present worldwide concern^1-3^. In addition to the lung damage traditionally associated with these infections, influenza virus can also cause or exacerbate cardiac dysfunction^4-10^. Ample evidence exists for the role of cardiac dysfunction in influenza-associated morbidity and mortality, including: (i) myocarditis is observed in a significant portion of hospitalized influenza patients^11-13^, (ii) heart damage at autopsy has been reported for fatal seasonal influenza cases^13-16^, (iii) severe cardiac damage was described in nearly all patients who died from infection with the 1918 pandemic influenza virus^17^, and (iv) cardiac events increase annually during flu season, especially among the unvaccinated^18,19^. Despite the implications for public health, little is known about the underlying mechanisms by which influenza virus causes heart pathology^11-16^.

Indeed, there is a debate within the clinical literature as to whether influenza virus directly or indirectly causes cardiac complications^6-11^. Although live virus has been detected in human and non-human primate heart samples, direct infection of the heart has rarely been investigated^20-24^. Instead, current dogma states that severely infected lungs produce a cytokine storm with systemic cardiotoxic inflammation, which indirectly drives cardiac dysfunction^6-11,25^. Attempts to resolve this fundamental question have been hindered by the lack of tractable animal model systems for influenza-mediated cardiac pathology^26,27^.

Laboratory mouse strains generally show minimal cardiac dysfunction upon influenza virus infection, even with high doses of virus^26-28^. Overcoming this obstacle, we recently reported that mice lacking the interferon-induced transmembrane protein 3 (IFITM3) suffer from severe cardiac electrical dysfunction and fibrosis upon influenza virus infection, thus providing a long-sought model for influenza-associated cardiac complications ^29^. IFITM3 is a protein involved in innate immunity that blocks the fusion of viruses with cell membranes, and deficiencies in IFITM3 are among the only known genetic risk factors in humans for developing severe influenza^30-36^. We observed that severe influenza virus-induced cardiac pathology in IFITM3 KO mice correlates with drastically increased and sustained viral loads in heart tissue when compared with rapid virus clearance in WT mice^29^. These results suggested a direct role for influenza virus replication in the heart in driving cardiac dysfunction, but the observed cardiac phenomena could not be decoupled from the severely heightened lung infection and inflammation that also occurs in IFITM3 KO mice^29,32,37^.

Here we sought to address the fundamental question of whether severe lung infection is sufficient to drive influenza-associated cardiac dysfunction, or whether replication in heart cells is required. We designed, rescued, and validated a recombinant influenza virus that is attenuated for replication in cardiomyocytes while being fully replication-competent and inflammatory in the lungs. Using this novel tool, we find that severe lung inflammation during influenza virus infection, even in highly infected IFITM3 KO mice, is not sufficient to drive cardiac dysfunction in the absence of virus replication in cardiomyocytes. Thus, direct infection and replication of influenza virus in cardiomyocytes is a primary determinant of cardiac pathology associated with severe influenza.

## Results

### Generation of influenza virus with cardiomyocyte-specific attenuation

Influenza virus strain A/Puerto Rico/8/1934 (H1N1), commonly known as PR8, is a pathogenic mouse-adapted virus that we previously showed disseminates from the lungs to the hearts of WT and IFITM3 KO mice^29^. As such, we chose this virus for a heart-specific attenuation strategy. Using reverse genetics techniques, we inserted into the influenza virus nucleoprotein (NP) gene segment, two copies each of target sequences for miR133b and miR206, two miRNAs that are expressed specifically in muscle cells, including cardiomyocytes (Fig 1A)^38-44^. Following miRNA target site insertions, we added a duplicated NP packaging sequence flanking the inserted target sequence. We rescued this novel recombinant virus in cell culture, and herein refer to it as PR8-miR133b/206. A control virus (PR8-miRctrl) containing a length-matched non-targeted sequence was described previously (Fig 1A)^38^. Both engineered viruses grew to similar high titers (>10^7^ TCID50/mL) in embryonated chicken eggs, indicating that relative replicative capacities of the engineered viruses were unaffected in the absence of specific miRNA targeting.

**Figure 1:**
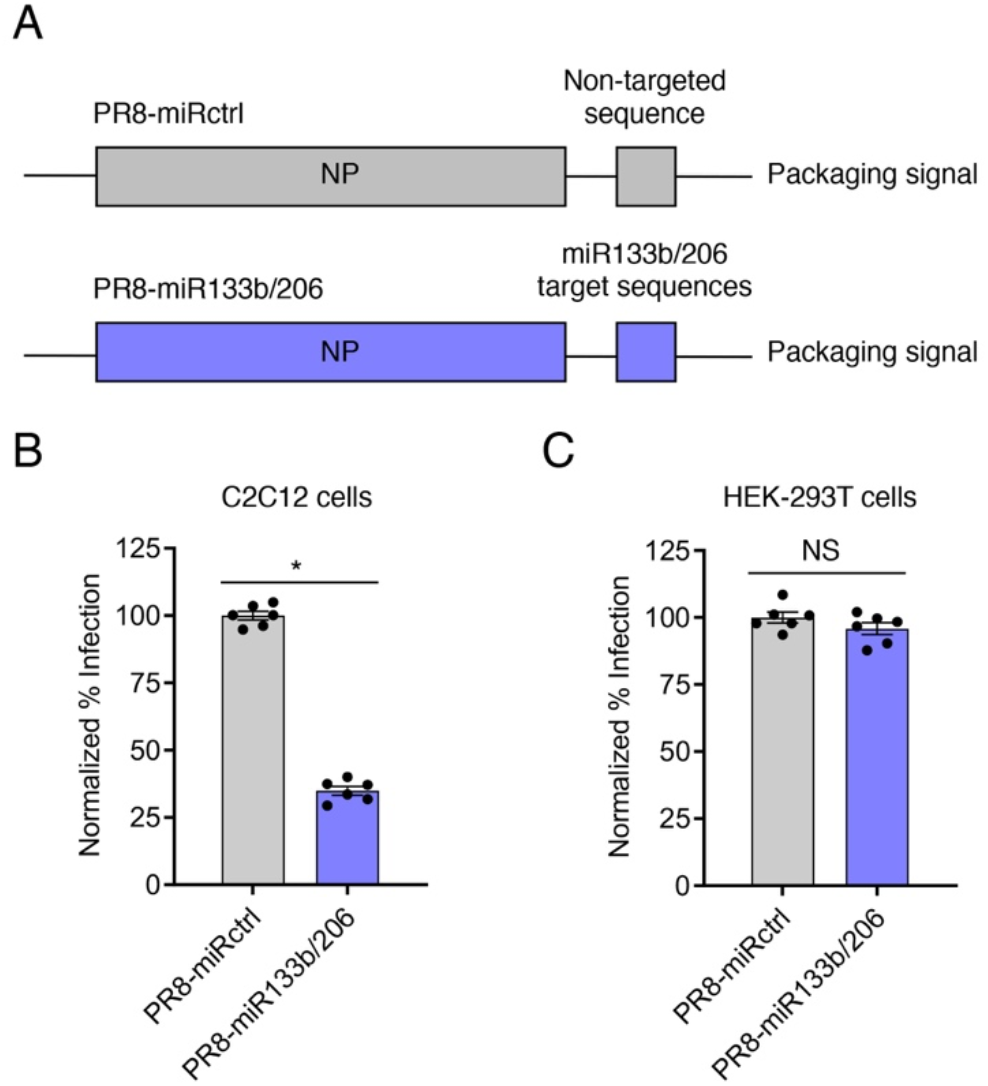
PR8-miR133b/206 is attenuated in myoblast cells *in vitro*. (A) Schematic of miRNA targeting strategy. Target sequences of two miRNAs expressed in cardiac cells, miR133b and miR206, or a length-matched random sequence were inserted into the influenza A PR8 NP gene, along with a duplicated NP packaging sequence, to generate replication competent virus with heart-specific attenuation (PR8-miR133b/206) or control virus (PR8-miRctrl). (B) C2C12 cells or (C) HEK-293T cells were infected with PR8-miR133b/206 or PR8-miRctrl for 24 h at an MOI of 2.5, and percent infection was determined by flow cytometry. Graphs represent normalized infection values. * p < 0.05 by unpaired t test; NS, not significant.

To validate that PR8-miR133b/206 is attenuated in cells expressing the relevant miRNAs, we infected a mouse myoblast cell line known as C2C12. Compared to the control virus, PR8-miR133b/206 was significantly attenuated in C2C12 cells, suggesting that targeting by miRNAs 133b and 206 potently restricts infection of myoblasts (Fig 1B). As a control, we observed no significant difference in infection by the two viruses in human HEK293T cells which do not express murine miR133b/206 (Fig 1C). Overall, we established PR8-miR133b/206 as an infectious, replication-competent virus that is attenuated in myocyte-like cells.

### Cardiomyocyte-specific miRNA-targeting of influenza virus prolongs survival of IFITM3 KO mice

To measure the overall pathogenicity of PR8-miR133b/206 compared to control virus, we infected WT and IFITM3 KO mice, and tracked their weight loss and survival. Consistent with enhanced disease severity in IFITM3 KO mice, the KO animals lost significantly more weight than WT mice in infections with both viruses (Fig 2A). Comparing the viruses within the individual mouse genotypes, we found that miR133b/206 targeting did not significantly alter the ability of influenza virus to induce weight loss (Fig 2A). Since weight loss during influenza virus infection is generally driven by cytokine-induced inappetence^45-47^, these data suggest similar levels of lung-derived inflammation were induced by the two viruses (examined below).

**Figure 2:**
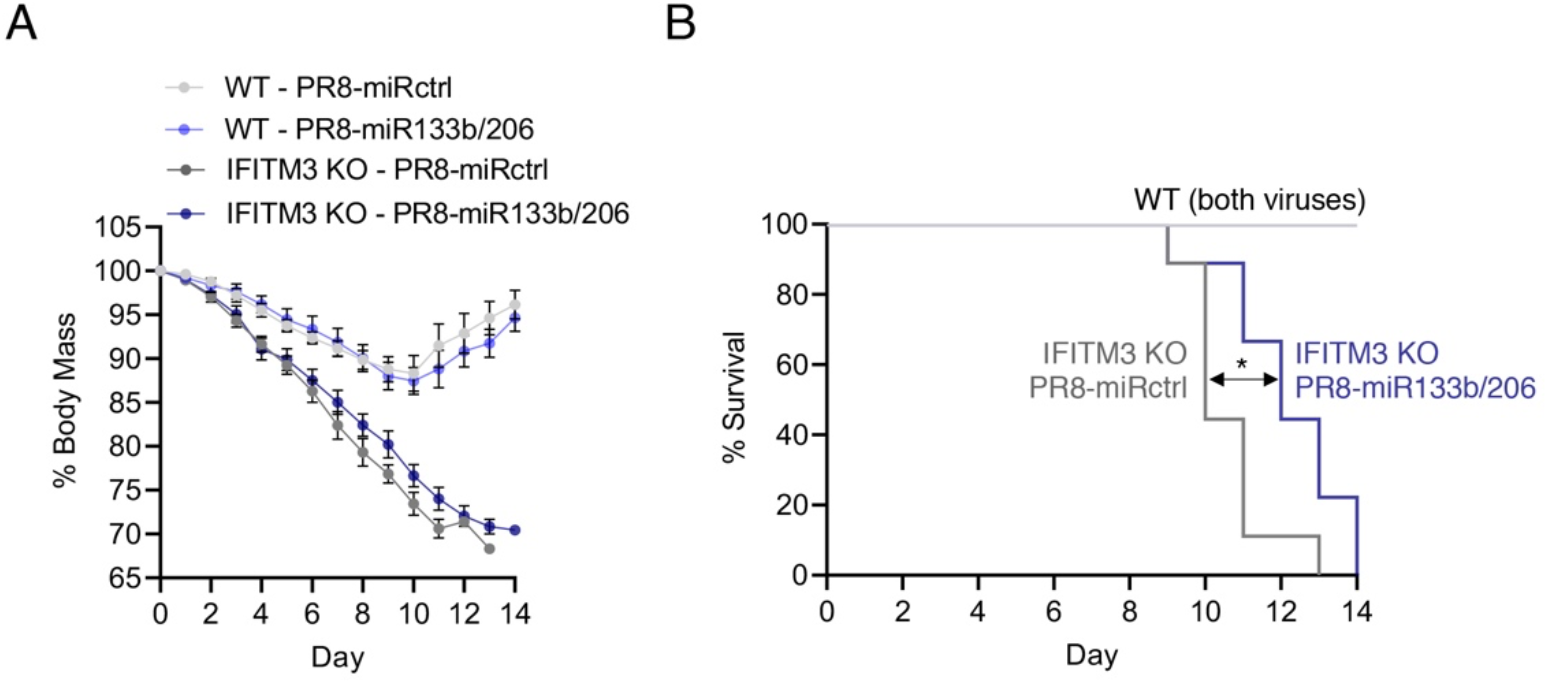
Cardiac infection with influenza virus decreases mean survival time in IFITM3 KO mice. WT and IFITM3 KO mice were intranasally infected with PR8-miR133b/206 or PR8-miRctrl (50 TCID_50_) and monitored daily for **(A)** weight loss and **(B)** survival. **A)** Points depict mean values collected from at least 3 experiments, and error bars represent standard deviation of the mean. Differences between WT and KO mouse weights were significant from day 4 onward with p < 0.05 by ANOVA with Tukey’s multiple comparisons test. Differences in weight loss when comparing PR8-miRctrl and PR8-miR133b/206 within the individual mouse genotypes were not significant. (B) Survival curves. The indicated p value is for statistical comparison of the IFITM3 KO survival curves (shown by double arrow) as calculated using a Gehan-Breslow-Wilcoxon test.

Similarly, all WT mice recovered from infections with either virus strain, while both infections were lethal in IFITM3 KO mice (Fig 2B). Despite similar weight loss in IFITM3 KO mice, infection with PR8-miR133b/206 resulted in a modest, statistically significant benefit in terms of survival as compared to infection with PR8-miRcontrol (median survival times of 10 d for PR8-miRctrl versus 12 d for PR8-miR-133b/206) (Fig 2B). These data suggest that viral replication in cardiomyocytes may contribute to more rapid lethality in IFITM3 KO mice, but, not surprisingly, that cardiomyocyte infection is not the sole cause of death. Overall, these outcome data are consistent with the premise that our recombinant viruses can decouple the impact of lung inflammation from the development of influenza-associated cardiac dysfunction.

### PR8-miR133b/206 is specifically attenuated in the heart *in vivo*

To further validate the utility of our engineered viruses in dissecting determinants for cardiac dysfunction, we first examined viral loads and lung inflammation after infection by PR8-miR-133b/206 and PR8-miRctrl. As shown in Fig. 3A, lung replication of PR8-miR133b/206 was comparable to that of PR8-miRctrl at both day 5 and day 10 post infection in WT mice. As expected, viral titers were higher in IFITM3 KO mice than WT mice (Fig 3A), but again were similar in the lungs when comparing PR8-miRctrl versus PR8-miR133b/206 (Fig 3A). To further confirm that PR8-miR133b/206 lung infections were not attenuated, we measured IFN and IL-6 levels, and found no significant difference for these proinflammatory cytokines in the lungs of mice infected by control or miR-targeted virus (Fig 3B,C). We also observed that IFITM3 KO mice, as expected, showed more severe histopathology than WT mice (Fig 3D). Indeed, both viruses induced cellular consolidation of the airways at indistinguishable levels (Fig 3E). Coupled with outcome data, the comparable lung pathology and inflammation demonstrate that the novel PR8-miR133b/206 virus is not attenuated in the lungs.

**Figure 3:**
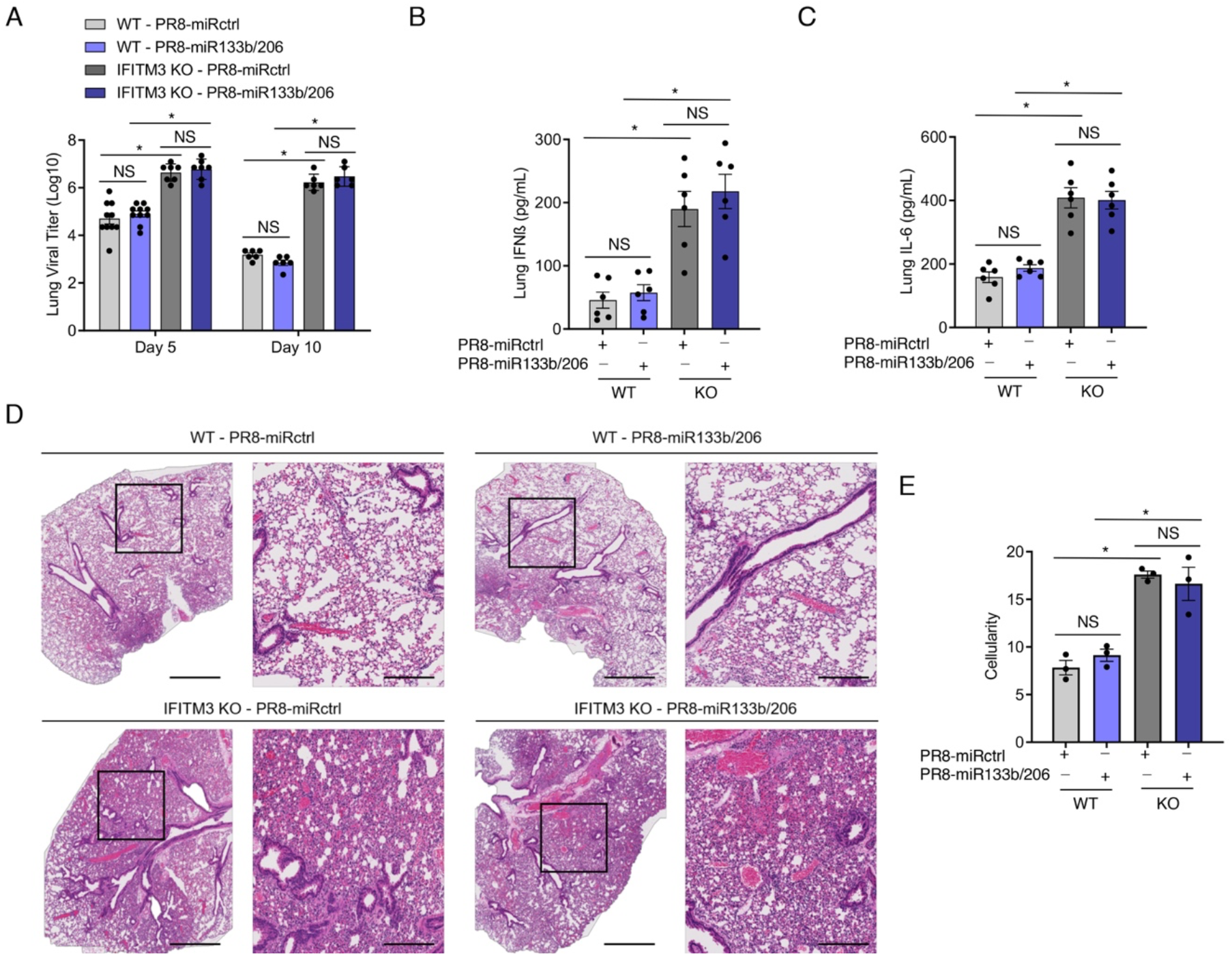
PR8-miR133b/206 is fully pathogenic in the lungs *in vivo*. WT and IFITM3 KO mice were intranasally infected with PR8-miR133b/206 or PR8-miRctrl (dose 50 TCID_50_). **(A-C)** Mice were euthanized on day 5 or 10 post-infection for measurement of virus titers **(A)** or ELISA quantification of IFNβ **(B)** and IL-6 **(C)** in the heart. Data points represent individual mice and bars represent mean values. Error bars depict standard deviation of the mean. Data points are from 3 independent experiments. Comparisons were analyzed by ANOVA followed by Tukey’s post-hoc test. * p < 0.05. **(D)** Mice were euthanized on day 10 post-infection for histological analysis of lung pathology. Boxed regions in the left image correspond to the magnified area depicted in the right image for each group. Scale bars represent 1mm and 400um for the left and right images, respectively. **(E)** Whole lung images as in D were quantified for overall cellularity using ImageJ. Data points represent individual mouse lung images and bars represent mean values. Error bars depict standard deviation of the mean. Comparisons were analyzed by ANOVA followed by Tukey’s post-hoc test. * p < 0.05.

Given the inclusion of muscle/cardiomyocyte-specific miRNA targeting sequences, we predicted that PR8-miR133b/206 infections would be attenuated in cardiac tissues. To test this prediction, we quantified virus titers in the same experimental mice used to derive the data from lungs (5 and 10 d post infection). As expected for PR8-miRctrl, we observed primarily low or undetectable viral titers in WT mouse hearts, and significantly higher titers in the cardiac tissue from IFITM3 KO mice at both timepoints (Fig 4A). In PR8-miR133b/206 samples, live virus was undetectable in WT hearts, and the mean titers were significantly lower in IFITM3 KO hearts compared to control virus, confirming overall attenuation of replication in the heart for the miR-targeted virus (Fig 4A). Cardiomyocyte-specific attenuation of the virus via miR targeting revealed an important role for the direct infection of cardiomyocytes in influenza-associated cardiac inflammation, as manifested in IFITM3 KO mice by: (i) a roughly 1-1.5 log decrease in mean cardiac viral titers, (ii) markedly reduced levels of inflammatory cytokines IFNβ and IL-6 (Fig 4B,C), and (iii) attenuated CD45-positive immune cell infiltration (Fig 3D,E).

**Figure 4:**
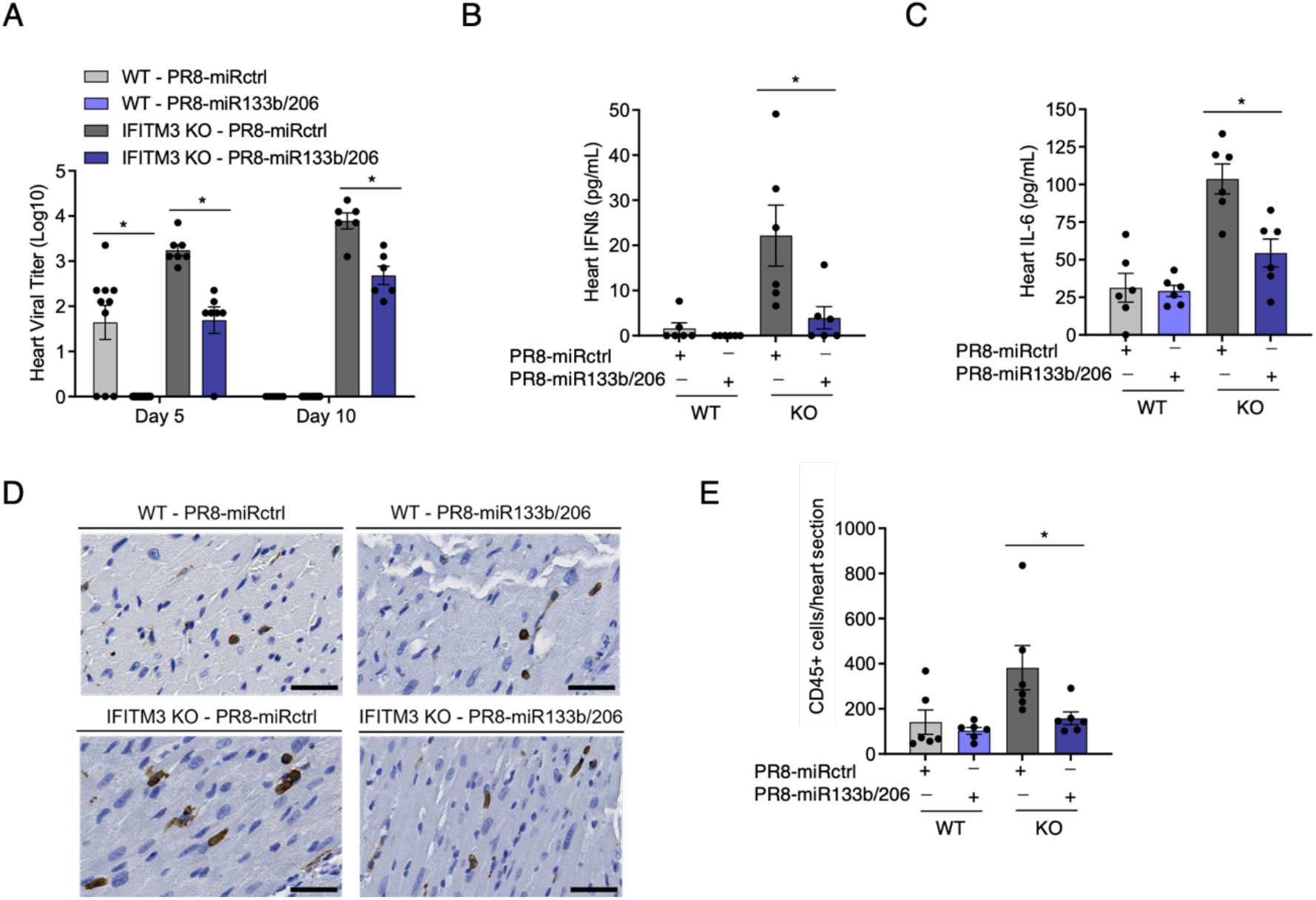
PR8-miR133b/206 is attenuated in the heart *in vivo*. WT and IFITM3 KO mice were intranasally infected with PR8-miR133b/206 or PR8-miRctrl (dose 50 TCID_50_). **(A-C)** Mice were euthanized on day 5 or 10 post-infection for TCID50 measurement of virus titers **(D)** or ELISA quantification of IFNβ **(B)** and IL-6 **(C)** and in the heart. Data points represent individual mice and bars represent mean values. Error bars depict standard deviation of the mean. Data points are from 3 independent experiments. Statistical comparisons were analyzed by ANOVA followed by Tukey’s post-hoc test. *p < 0.05. **(D)** Mice were euthanized on day 10 post-infection for histological analysis of CD45+ immune cell infiltration in the heart. Images shown depict areas of immune cell infiltration indicated by brown staining. Scale bars represent 50um. **(E)** Whole heart images were quantified for CD45+ cells using ImageJ. Data points represent individual mouse heart images and bars represent mean values. Error bars depict standard deviation of the mean. Comparisons were analyzed by ANOVA followed by Tukey’s post-hoc test. * p < 0.05.

Overall, we have established a controlled experimental system to interrogate roles of lung inflammation versus direct cardiac infection in influenza-associated cardiac dysfunction. Namely, IFITM3 KO mice infected with PR8-miRctrl or PR8-miR133b/206 allow direct comparison of heart phenotypes in animals with equivalently severe lung infections, but with or without high virus replication in the heart.

### Virus replication in the heart is required for robust induction of cardiac fibrosis and electrical dysfunction

Fibrosis is a broadly observed consequence of severe infectious insults to cardiac tissue in humans. Because IFITM3 KO mice exhibit significant cardiac fibrosis following infection with influenza virus^29^, and because lung inflammation is not attenuated for our heart-attenuated virus, we could test for the first time whether cardiac pathology results directly from heart infection or is induced indirectly by severe lung inflammation. We thus collected hearts from WT and IFITM3 KO mice at day 10 post-infection, and first performed histological analysis of fibrosis using Masson’s Trichrome staining. Examination of heart sections revealed blue-stained fibrotic lesions that were most apparent in hearts from IFITM3 KO mice infected with PR8-miRctrl (Fig 5A). Indeed, fibrotic lesions in WT samples and those from IFITM3 KO mice infected with PR8-miR133b/206 were minimal (Fig 5A). Quantitative analysis of images from multiple mice confirmed that fibrotic staining in heart samples from IFITM3 KO mice infected with miR-targeted virus was significantly decreased compared to infection with miRctrl virus (Fig 5B). Thus, attenuation of virus replication in the heart correlates with less cardiac fibrosis following infection, indicating that direct virus replication in cardiomyocytes is required for development of influenza-associated cardiac fibrosis.

**Figure 5:**
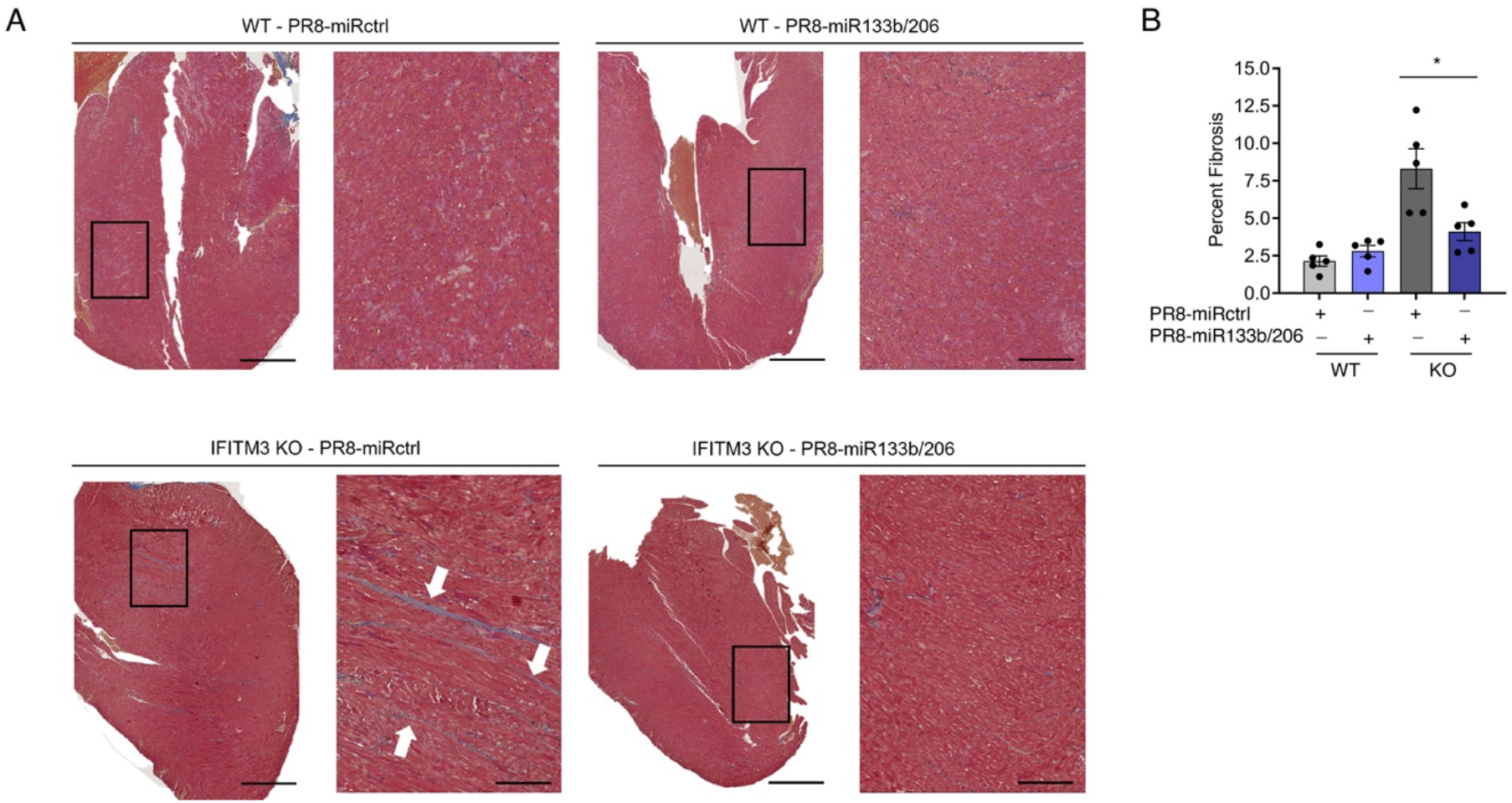
Virus replication in the heart is necessary to induce fibrosis during infection. WT and IFITM3 KO mice were intranasally infected with PR8-miR133b/206 or PR8-miRctrl (50 TCID_50_). **(A)** Hearts were collected on day 10 post-infection, and sections were stained with Masson’s trichrome stain, in which blue staining is indicative of fibrotic collagen deposition. Histological processing and image acquisition were performed by the OSU Comparative Pathology and Mouse Phenotyping Core Facility on heart tissue samples provided by Adam Kenney. A representative heart section is shown for each genotype-virus combination. Boxed areas are regions magnified in the far-right images. **(B)** Percent fibrosis was calculated by quantifying ratio of blue pixel intensity to total pixel intensity for each heart section. Each point represents a heart from an individual mouse, and bars represent mean values. Error bars represent standard deviation of the mean. Comparisons were analyzed by ANOVA followed by Tukey’s post-hoc test. *p < 0.05.

Given that cardiac fibrosis is a well-established risk-factor for cardiac arrythmia^48,49^, and that fibrosis was significantly decreased in infection with PR8-miR133b/206 (Fig 5), we tested whether cardiac electrical dysfunction induced by influenza virus similarly requires direct infection of cardiomyocytes. We performed electrocardiogram (ECG) measurements on WT and IFITM3 KO mice before infection and at several timepoints after infection with PR8-miRctrl or PR8-miR133b/206. While cardiac function in WT mice was largely unchanged throughout infection, IFITM3 KO mice showed depressed heart rates and increased RR (interbeat) interval averages during the course of infection with PR8-miRctrl (Fig 6A). We also observed irregular ECG tracings in these KO animals infected with PR8-miRctrl, indicative of an abnormally arrhythmic heartbeat (Fig 6B). Remarkably, an arrhythmic phenotype was largely absent in the majority of IFITM3 KO mice infected with PR8-miR133b/206 (Fig 6C). Specifically, RR interval ranges (defined as the longest RR interval minus the shortest RR interval)^29^ calculated from the entirety of our ECG readings in multiple mice were significantly lower in KO mice infected with PR8-miR133b/206 compared to PR8miR-ctrl. Importantly, these latter data indicate an attenuation of pathological arrhythmic heart dysfunction when virus replication in the heart was decreased (Fig 6C). Overall, the attenuation in heart viral titers observed for infections with PR8-miR133b/206 is accompanied by decreased cardiac electrical dysfunction, despite the robust virus replication and inflammation in the lungs. We conclude that influenza-associated cardiac fibrosis and electrical dysfunction requires direct virus replication in the heart.

**Figure 6:**
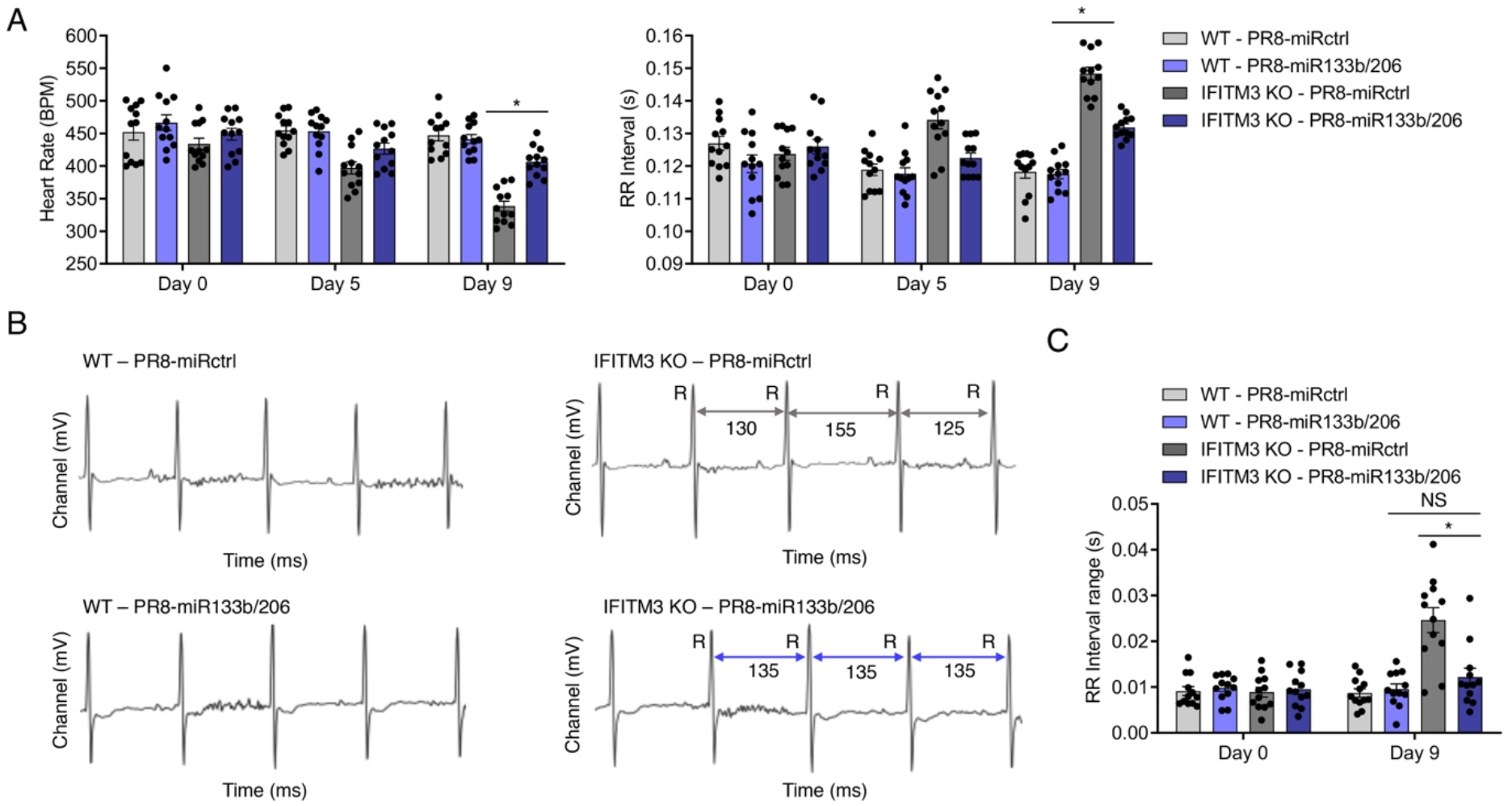
Virus replication in the heart drives cardiac dysfunction during infection. WT and IFITM3 KO mice were intranasally infected with PR8-miR133b/206 or PR8-miRctrl (50 TCID_50_). **(A)** ECG measurements over the time course of infection. Data were collected over at least 3 independent experiments. Each point represents an individual mouse, and bars represent mean values. Error bars represent standard deviation of the mean. Comparisons were analyzed by ANOVA followed by Tukey’s post-hoc test. *p < 0.05. **(B)** Example ECG readings from each genotype-virus combination. Selected RR intervals of the infected KO mice are highlighted by grey (PR8-miRctrl) or purple (PR8-miR133b/206) double arrows. **(C)** RR interval ranges, defined as the difference between the longest and shortest RR intervals over an ECG measurement period of 5 minutes, were calculated for individual mice on day 9 post-infection. Each point represents an individual mouse, and bars represent mean values. Error bars represent standard deviation of the mean. Comparisons were analyzed by ANOVA followed by Tukey’s post-hoc test. *p < 0.05.

### Rat and human cardiomyocytes are susceptible to influenza virus infection

To date, direct infection of mammalian cardiac cells by influenza virus has been rarely investigated, but would provide important support for an infection-dysfunction link in pathology. To confirm that primary rodent cardiomyocytes are permissive to influenza virus infection, we purified rat neonatal cardiomyocytes and infected these cells for 24 h with increasing MOIs of PR8 expressing GFP (PR8-GFP). As shown in Fig. 7A, imaging of GFP fluorescence revealed that the cardiomyocytes were infected by influenza virus (Fig 7A). We next tested whether human cardiomyocytes are able to be infected. For this, we utilized human induced pluripotent stem cell-derived cardiomyocytes, derived from two independent sources and culture methods. Similarly to the rat cells, we readily visualized PR8-GFP infection of Fujifilm iCell cardiomyocytes, as indicated by GFP expression at 24 h post infection (Fig 7B). In addition, cardiomyocytes cultures differentiated in-house from human stem cells were infected by PR8-GFP at similarly high levels (Fig 7C). Notably, GFP-positive cardiomyocytes that were previously beating in these cultures ceased to beat by 48 h post infection (Supp Movie 1), while mock-infected wells continued to be populated by contractile cells (Supp Movie 2). At a mechanistic level, we found that influenza virus infection of the cardiomyocytes significantly increased cleaved caspase 3 levels, indicating activation of cell death pathways (Fig 7D,E). Taken together, we conclude from these experiments that cardiomyocytes are susceptible to direct influenza virus infection, which activates cardiomyocyte death pathways.

**Figure 7:**
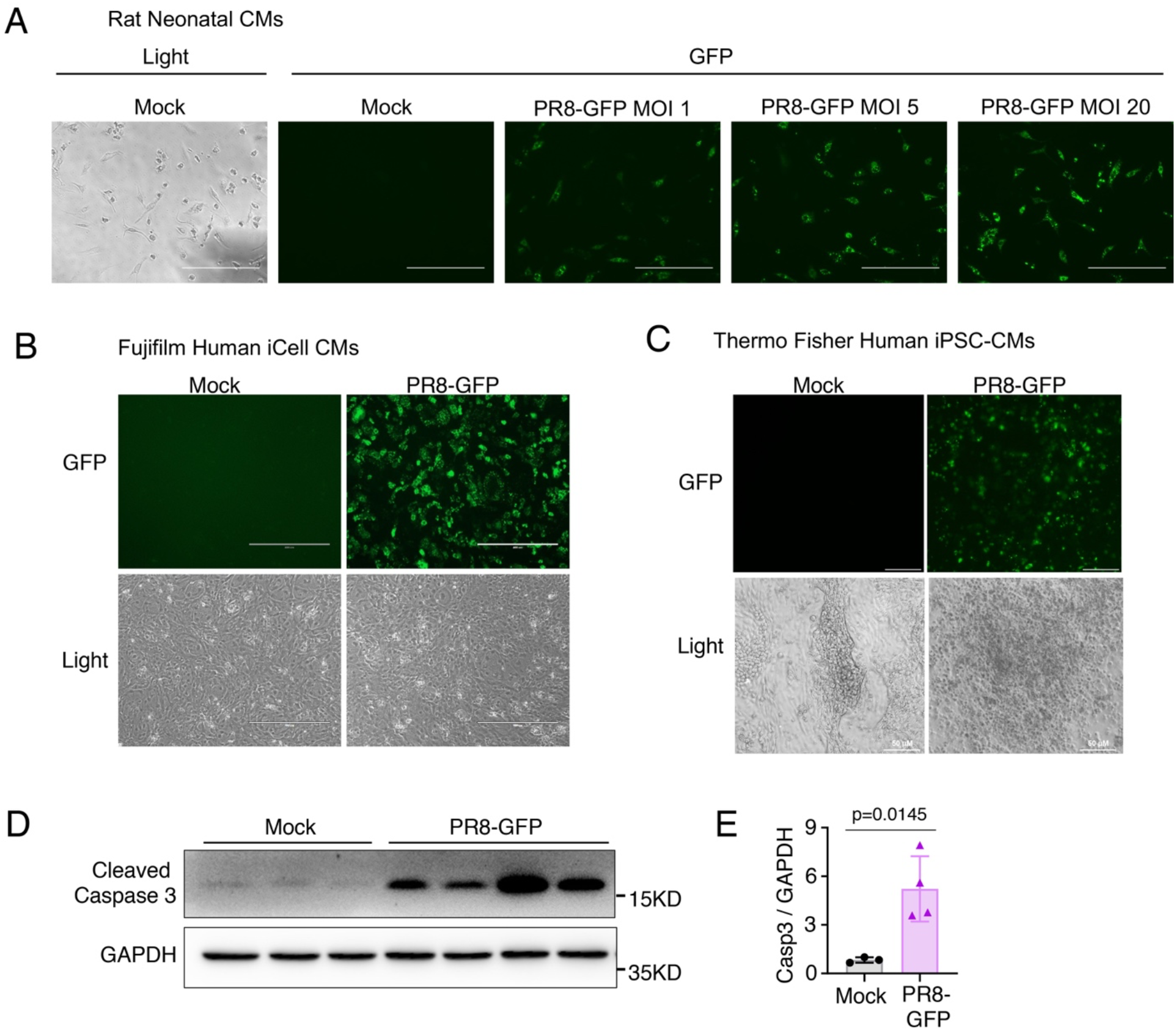
Primary cardiomyocytes are susceptible to influenza virus infection. **(A)** Cardiomyocytes (CMs) purified from neonatal rat hearts, subjected to influenza virus strain PR8-GFP infection for 24 h at the indicated MOIs, and imaged by fluorescent (GFP) and light microscopy. Scale bars, 400 um. **(B)** Human iCell CMs were purchased from Fuji Film, cultured, infected with PR8-GFP (MOI 5) for 48 h, and imaged by fluorescent (GFP) and light microscopy. **(C)** Human CMs differentiated from Thermo Fisher-purchased induced pluipotent stem cells (iPSC) were infected with PR8-GFP (MOI 5) for 48 h, and imaged by fluorescent (GFP) and light microscopy. **(D)** Samples from 3 independent experiments as in **C** were blotted for cleaved caspase-3 and GAPDH. The two rightmost lanes are replicate infected samples from the same experiment. **(E)** Quantification of Caspase-3 (Casp3) levels in **D** normalized to GAPDH levels.

## Discussion

Cardiac manifestations of influenza virus infection are widely attributed to severe lung inflammation, which contributes to systemic tissue damage and exacerbates pre-existing heart conditions^50-52^. However, given that we lack non-invasive clinical tests to identify direct heart infection by influenza virus in living humans, it has been difficult to determine the relative contributions of lung inflammation versus direct virus-induced damage to the heart in cardiac dysfunction during severe infection. To address this fundamental question, we turned to an animal model, specifically, IFITM3 KO mice, which serve as a severe infection model to study influenza-induced cardiac dysfunction^29^. IFITM3 alters membrane properties to disfavor virus-to-cell fusion^53-57^. IFITM3 KO mice thus experience increased cellular infection and spread in lungs, spleen, and heart^29^, organs that are naturally susceptible to infection in WT mice. Importantly, viremic infection of other organs, such as the brain, liver, or kidneys, is not observed in IFITM3 KO mice^29^, thus providing a severe infection model recapitulating the tissue-specific distribution of influenza virus dissemination. Further supporting its relevance in dissecting influenza pathologies, genetic deficiencies in IFITM3 associate with susceptibility to severe disease in humans^30-36^.

To manipulate the ability of influenza virus to replicate in cardiomyocytes, we generated a novel recombinant virus strain with cardiomyocyte-specific miRNA target sites. We found that insertion of microRNA target sites for miR133b and 206 into the NP genome segment of IAV strain PR8 effectively attenuated infection specifically in murine myocyte-like cells *in vitro* and in the heart *in vivo* (Fig 1, 4). The reduction of virus load in the heart correlated with less severe cardiac fibrosis, inflammation, and electrical dysfunction, though lung virus replication and inflammation remained robust and comparable to control virus (Fig 3-6). Thus, we identified that direct infection of heart cells is required for cardiac dysfunction during influenza virus infection. Our findings overturn the notion that severe lung inflammation is sufficient for influenza-associated cardiac pathologies. Since cardiac complications of severe influenza are often seen in hospitalized patients^11-13^, our results may suggest that direct infection of the human heart is more common than currently appreciated.

Both cardiac fibrosis and cardiac electric dysfunction were reduced in IFITM3 KO mice infected with PR8-miR133b/206, despite retention of residual virus in the heart (Fig 4-6). These data suggest that a threshold of virus in heart tissue is tolerated without producing significant pathology. This notion is strengthened by the fact that WT mice often have low, quickly cleared, levels of virus in the heart at early timepoints post-infection, but rarely exhibit significant cardiac dysfunction^27-29^. Alternatively, infection of cell types in the heart, in addition to cardiomyocytes, may occur without major pathological outcomes. Importantly, however, we observed no protective advantage of reduced cardiac infection in IFITM3 KO mice in terms of weight loss (Fig 2A), a finding that underscores the severity of the lung infection experienced by IFITM3 KO mice.

Several key issues remain to be addressed by future approaches in dissecting cardiac pathogenesis of influenza virus. Of particular interest is the mechanism by which the virus spreads from the primary site of infection (respiratory tract/lungs) to the heart and how productive infection in the heart is achieved thereafter. Single cell analyses may prove useful for identifying cell subsets in the heart that are infected initially by influenza virus. Additionally, virus strain-specific differences influencing not only overall virulence, but also tissue tropism, may influence cardiac infection and pathology. Indeed, certain strains of influenza virus have been shown to preferentially infect upper vs lower respiratory tract and other sites of extrapulmonary tropism are noted for particular virus strains^58-62^. Identifying viral factors that influence cardiac infection could prove crucial for predicting and treating cardiac manifestations of both circulating and emerging viruses. Finally, there is much to be learned about the clinical role of cardiac infection in humans, particularly in individuals with deleterious IFITM3 SNPs, who may have a greater risk for direct influenza virus infection of the heart and cardiac pathology. Overall, extrapulmonary manifestations of respiratory virus infections are increasingly appreciated as important aspects of disease that will require continued research. Understanding the direct and indirect effects of virus in extrapulmonary tissues, such as the direct effects of influenza virus on the heart as uncovered here, will be critical for combating these noncanonical disease pathologies.

## Materials and Methods

### Virus generation, propagation and titering

PR8-miRctrl and PR8-miR133b/206 were generated as previously described^38-44^. Briefly, two copies each of target sequences for miR133b and miR206, or a length-matched untargeted sequence, were cloned and ligated into the 3′ UTR of the NP gene along with a duplicated 3’ NP packaging sequence. The recombinant viruses with modified NP segments were rescued using reverse genetic techniques and plaque purified. PR8-GFP was previously described^63,64^.

Viruses were propagated in 10-day-old embryonated chicken eggs (Charles River Laboratories) for 48 hours at 37°C and titered on MDCK cells. For determining organ titers, tissues were collected and homogenized in 500ul of PBS, flash-frozen, and stored at -80°C prior to titering on MDCK cells.

### Mouse infections

Mice eight to12 weeks of age were anesthetized with isoflurane (Henry Schein Animal Health) and intranasally infected with PR8-miRctrl or PR8-miR133b/206 (50 TCID50) in sterile saline. Mice were monitored daily for weight loss and morbidity, and sacrificed if weight loss exceeded 30% of starting body mass or other endpoint criteria (severe hunched posture, lack of ambulation) were met. All procedures were approved by the OSU IACUC.

### Cell lines, cell line infections, and flow cytometry

C2C12, HEK-293T, and MDCK cells were grown in Dulbecco’s Modified Eagle’s Medium (DMEM) supplemented with 10% Equafetal bovine serum (FBS; Atlas Biologicals) at 37°C with 5% CO2 in a humidified incubator. C2C12 and HEK-293T cells were infected with PR8-miRctrl or PR8-miR133b/206 at an MOI of 1.0 for 24 hours. For determination of IAV-infection percentages via flow cytometry, cells were stained with anti-H1N1 IAV NP (BEI resources, clone 4F2) and Alexa488-conjugated secondary antibody (Thermo Scientific). Flow cytometry was performed on a FACSCanto II flow cytometer (BD Biosciences) and analyzed using FlowJo software.

### Primary cardiomyocytes

Rat neonatal cardiomyocytes were prepared as described previously^65^. Fujifilm iCell cardiomyocytes were purchased from FujiFilm Cellular Dynamics Inc. (FCDI) (cat# C1006) and cultured in hiPSC-CM plating medium (FCDI, Cat# M1001) for 2 d followed by maintenance medium (FCDI, cat# M1003) for 10 d. In-house differentiated primary cardiomyocytes were cultured as follows. Human iPSCs were purchased from Thermo Fisher (cat.# A18945,), and cultured on Matrigel-coated plates (Corning, cat# 356231) using StemFlex medium (Thermo Fisher, cat.# A3349401) as described previously^66^. Cardiac differentiation was initiated by using a small molecule–based protocol^67^. PSCs (passage 20–35) were cultured until 80%–90% confluence, and their medium was replaced with cardiac differentiation basal medium consisting of RPMI (Thermo Fisher cat# 11875119), B27 supplement minus insulin (Theremo Fisher, cat.# A1895601). For the first two days, the basal medium was supplemented with 9 μM CHIR-99021 (Selleckchem, cat# S2924). The medium was then replaced with cardiac differentiation basal medium plus 5 μM IWR-1 (Sigma, cat# I0161). Differentiated cells were maintained in cardiac differentiation basal medium for 2 days, which was then replaced with cardiac proliferation medium consisting of RPMI, B27 supplement (Thermo Fisher, cat# 17504044) for another 4 days. Beating of iPSC-differentiated cardiomyocytes (hiPSC-CMs) was typically observed between days 8–12 of differentiation. Influenza virus infections were performed on cultured in which beating cardiomyocyte cell clusters could be visually observed. For all infections of primary cells, virus was added directly to culture media and infection was allowed to proceed for 24-48 h.

### ELISA

IFNβ and IL-6 concentrations in organ homogenates were analyzed using mouse DuoSet ELISA kits (R&D Systems).

### Electrocardiography

For subsurface electrocardiograph (ECG) recordings, anesthesia was provided by isoflurane in oxygen at a flow rate of 1.0 L/min. Mice were placed in a prone position on a heated pad to maintain body temperature, and subcutaneous electrodes were placed under the skin (lead II configuration). ECGs were recorded for five minutes on a Powerlab 4/30 (AD Instruments).

Anesthesia was maintained for the duration of the reading. ECG traces were analyzed using LabChart 8 Pro software (AD Instruments).

### Immunohistochemistry

For immunohistochemistry, hearts were fixed in 10% formalin and maintained at 4°C until embedded in paraffin. Hearts were sectioned by the OSU Comparative Pathology and Mouse Phenotyping Shared Resource. Masson’s trichrome staining was used to identify fibrotic replacement of cardiac tissue. Digital images of heart sections were generated using Aperio ImageScope software (Leica Biosystems). Images were analyzed via ImageJ (Version 2.0.0) as previously described ^29^.

## Author Contributions

Experiments were conceived by ADK, JSY, RAL, JM, MVSR, and CC. Experiments were performed by ADK, LZ, AZ, PJD, CC, HZ, AE, JK, LD, FA, and NVM. Design and rescue of miRNA-targeted viruses was performed by SA, CG, and RAL. Data was analyzed by ADK, MVSR, NK, CC, and JSY. The manuscript was written by ADK and JSY with editorial input from all authors.

## Acknowledgments

This research was supported by NIH Grants AI130110, AI151230, and AI142256 to JSY, grant AI146252 to MVSR, grants AI148669 and AI132962, grant HL154001 to JSY and FA, and grant AI146690 to JSY and MVSR. ADK was supported by The Ohio State University Systems and Integrative Biology Training Program funded by NIH Grant GM068412 and The Ohio State University Presidential Fellowship. AZ and AE were supported by an Ohio State University Infectious Diseases Institute training grant funded by NIH Grant AI112542 and The Ohio State University College of Medicine. AZ was also funded by the National Science Foundation GRFP Fellowship.

